# Emergent Effect of Mixed Crowders on Protein Dimerization

**DOI:** 10.1101/2025.04.27.650838

**Authors:** Rajendra Rath, Sweta Pradhan, Mithun Biswas

## Abstract

In vivo protein-protein association is modulated by non-specific interactions with the cellular matrix, known as crowding. Replicating the heterogeneity and complexity of cellular conditions in experiments and simulations to quantify the crowding influences remains a formidable challenge. As an alternative, homogeneous mixtures of polymers or proteins as crowders are employed, although they provide limited insight to the actual effect. In this work, we present a coarse-grained model of GB1 dimerization in lysozyme crowders by adjusting the GB1-lysozyme effective potential parameters obtained through high-resolution Martini simulation. Employing these parameters it is shown that GB1 dimerization is destabilized in presence of lysozyme, as observed in experiments. The model is further extended to incorporate binary crowder mixtures for crowder species of various sizes and interaction potentials. It is found that depending on mixing conditions and crowder volume fraction, dimerization can either be stabilized or destabilized. In particular, in presence of crowders of different sizes, the smaller one plays the dominant role. Finally, we observe that the overall change of stability due to crowding emerges from a delicate balance between enthalpy and entropy that cannot be predicted by considering the property of single crowder species. These results provide invaluable insights for interpreting experimental observations under cell-like conditions.

## 1 Introduction

A cellular environment consists of a diverse collection of macromolecules, DNA-RNA, osmolytes and metabolytes. Evidences suggest that all of these species together can occupy ~ 30-40% of cellular volume that interacts non-specifically with other macromolecules in the vicinity affecting key cellular events such as protein folding and structural dynamics, ^1–9^ association and assembly, ^1,10–17^ enzyme catalysis, ^18–20^ signal transduction, ^21–23^ diffusive dynamics ^24–29^ as well as phase separation. ^30–33^ To capture the effect of this environment, effort has been made to mimic macromolecular crowding by considering one single species of polymers (for example, Ficoll, dextran) or proteins. ^34–36^ However, a single species of crowders lack the heterogeneous interactions and conditions present *in vivo*. A heterogeneous solution of many macromolecules as obtained in a lysis buffer is closer to the conditions present in cell, although in presence of a large number of competing interactions the interpretation of crowding effects becomes challenging. ^37^ A binary mixture of two crowding species provides crucial insights regarding how crowding influences evolve with increasing heterogeneity in the environment in a systematic manner. ^38–41^

In this work we employ protein-protein association, a ubiquitous cellular event regulating a variety of biological processes, ^1,42^ to understand the role of mixed macromolecular crowding. Early studies of protein association employed polymeric crowders showing the influence of volume exclusion in crowder induced stabilization. ^12,43–48^ More recently the use of proteins as crowders showed that, in addition to excluded volume effect, soft interactions (such as, van der Waals, electrostatic, hydrophilic and hydrophobic interactions) also play a major role in deciding the net stabilization or destabilization of the process. ^14,34,49,50^ In particular, lysozyme has been shown to destabilize the dimer formation of the globular GB1 proteins through soft attractive interaction, whereas BSA can stabilize the association by soft repulsive interaction. ^14^ The polymeric crowder PEG stabilizes the coupled folding-binding of intrinsically disordered proteins (IDPs), steroid receptor coactivator 3 (ACTR) and the nuclear coactivator binding domain of CBP/p300 (NCBD), through excluded volume effect. ^36^ Cytochrome c - Cytochrome c peroxidase (Cc-CcP) complex is marginally stabilized in saccharides. ^51^ The binding affinity of histidine carrier protein HPr with its partner EIN is reduced in protein crowder BSA, but enhanced in polymeric crowder Ficoll-70. ^52^ Interestingly, PEG has been shown to have either small effect on certain protein associations (TEM-BLIP association) ^13^ or a larger effect (GB1 dimerization). ^34^

Computer simulations provide a versatile tool to study protein-protein association under crowded conditions. ^1,53–57^ All-atom molecular dynamics simulations of a cell-like environment (for example, E. Coli bacterial cell having ~ 100 M atoms) is computationally demanding. ^58–62^ To address relevant timescales for protein association, with a balance between system size and required resolution, coarse-graining (CG) of the protein-crowder structures and interaction potentials is often employed as an useful strategy with limited computational resources. ^15,41,56,63–66^ Residue-level CG models can incorporate the flexibility of proteins and crowders along with the solvent treated explicitly or implicitly to investigate the influence of a crowded environment on protein folding and association ^53^ or free energy landscape. ^15^ Further simplified CG models, for example where both the proteins and crowders are assumed as spheroids, lacks description of structural changes induced by crowding, although they are able to capture the influence of complex binary or multi-component mixtures on the free energy change. ^41,57^

A systematic investigation of the influence of a heterogeneous crowding environment on protein-protein association is lacking, despite having several studies of protein association in mixed crowder solutions,. ^39,41,67–70^ To this end, for the work presented here, homodimerization of a well-known model system, immunoglobulin-binding protein G of the B1 domain of the Streptococcus species, in presence of single crowders as well as binary mixtures of two crowder species, is investigated. GB1 dimerization in presence of various protein and polymeric crowders has been studied previously in detail. ^14^ Here we employ a basic CG representation to simulate GB1 dimerization under the influence of attractive or repulsive protein-crowder interaction potential. The binary crowder mixture is prepared by using combinations of crowders of two different sizes (that is, larger and smaller than the protein) and two different interaction potentials (that is, repulsive and attractive) in different mixing ratios. Thus the simulated binary solutions encompass the following crowding scenarios: 1) Two repulsive crowder species of different sizes, 2) One large repulsive crowder and one small attractive crowder species, 3) One large attractive crowder and one small repulsive crowder species, and 4) Two attractive crowder species of different sizes. To understand the free energy change in each case, the enthalpic and entropic contributions are evaluated, along with an overview of protein-crowder configurations populated in different mixing solutions. Our results indicate emergent effects in a binary mixture due to complex balance of enthalpic and entropic terms.

The article is organized as follows: In Section 2, we derive the protein-crowder effective potential and outline the calculation of change of free energy ΔΔ*F* from CG simulations. Section 3 presents the variation of ΔΔ*F* with *ϕ* for single and mixed crowder species along with a description of corresponding enthalpic and entropic contributions. Section 4 summarizes the main findings of the work with a future outlook.

## 2 Methods

### CG simulation details

In a cell-like environment, reacting protein species diffuse through the cytoplasm and form the product at close contact. To implement this diffusion-reaction dynamics, a particle-based reaction diffusion simulation model (ReaDDy) ^71–73^ is used. All protein and crowder species are modeled as spherical particles having certain hydrodynamic radii. The diffusion coefficient of a species is obtained from the Stokes-Einstein relation using the viscosity of water (*η*) and corresponding hydrodynamic radius. The diffusive dynamics of the particles follows the overdamped Langevin equation

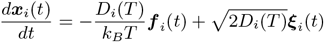

where ***x***_*i*_(*t*) is the position of the *i*-th particle at time *t, D*_*i*_(*T*) is the isotropic diffusion coefficient for *i*-th particle, *k*_*B*_ is the Boltzmann constant, *T* is the system temperature in K, ***f***_*i*_(*t*) denotes the deterministic force and ***ξ***_*i*_ are independent, Gaussian distributed noise with moments

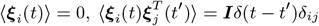

where ***I*** is the identity matrix.

In experiments, GB1 dimerization occurs in presence of attractive as well as repulsive crowders. ^14^ To study the destabilizing effect of attractive crowders on dimer stability, lysozyme is used as the crowding species having hydrodynamic radius *R*_*h*_ ~ 20 Å. ^74^ In a previous study, ^15^ attractive protein-crowder interaction was modeled employing the usual 12-6 Lennard-Jones (LJ) potential of the form,

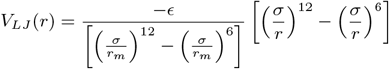

where *σ* is the distance where *V*_*LJ*_ (*r*) = 0, *r*_*m*_ is the distance at which *V*_*LJ*_ (*r*) reaches minimum and *ϵ* is the well-depth. However, the model predicted stabilization of GB1 dimerization in presence of lysozyme crowders, in contrary to experiments. ^14^ To improve on the effective potential of mean force (PMF) between lysozyme and GB1, here coarse grained molecular dynamics simulation of one GB1 molecule in presence of one lysozyme was performed using Martini force field ^65^ (see Supporting Information for details). It is found that the GB1-lysozyme PMF is much narrower compared to a 12-6 LJ potential and can be fitted to a generalized LJ-type potential with exponents *m* = 48 and *n* = 28 (See Supporting Information (Fig. S1)). To fit the LJ-type potential to the simulation data, it is further noted that during MD simulations GB1 and lysozyme adopt conformations which are far from globular and the obtained PMF is averaged over all possible orientations of these conformations. Hence, the PMF derived from MD simulations was shifted in such a way that the PMF is zero at contact distance between GB1 and lysozyme (*ϕ*(*r*_contact_) = 0, where *r*_contact_ is the sum of hydrodynamic radii of GB1 and lysozyme). It is also found that LJ-type potential is analogous to modified LJ (Kim-Mittal (KM)) potential employed in previous studies. ^56^ A comparison between different LJ potential forms is shown in Fig. 1. Since the depth of the LJ-like potential (*ϵ*) still remains undetermined, we analyze the influence of varying potential depth on crowder induced stabilization in the next section.

**Figure 1:**
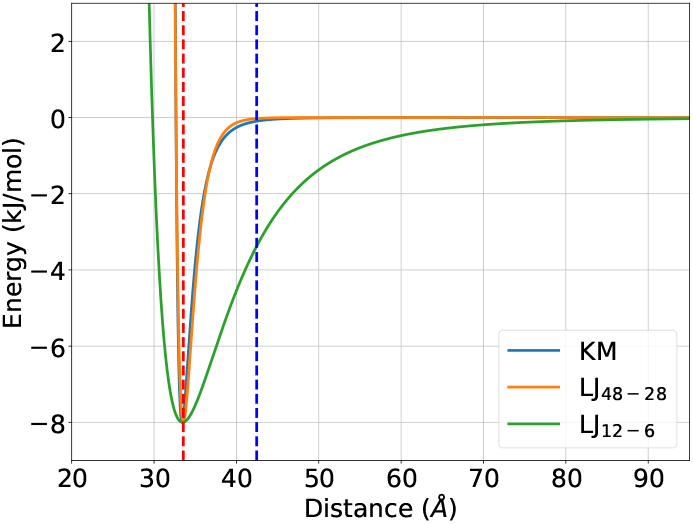
Comparison of Lennard-Jones potentials: 12-6 LJ (*green*), 48-28 LJ (*orange*), KM model (*blue*). The dotted red and blue traces indicate the minima of the LJ potentials and the cutoff distance for the 48-28 LJ potential, respectively.

In addition to large macromolecular crowders having strong but nonspecific interaction with the reacting proteins, there can be many small but weakly interacting macromolecules in the cellular matrix which stick to the protein and may change the association free energy. ^37,75^ To investigate this scenario, we introduced a smaller crowding species (*R*_*h*_ ~ 12 Å) compared to the protein monomers (*R*_*h*_ ~ 13.5 Å). The exponents *m* − *n* for the LJ-type potential describing the protein-small (12 Å) crowder attractive interaction is found by fitting to the KM model as detailed in the Supporting Information (Fig. S4).

**Table 1:** Parameters for ReaDDy simulation.

| Parameter | Value |
| --- | --- |
| Box size (Å<sup>3</sup>), $b$ | (150)<sup>3</sup> |
| Temperature(K), $T$ | 300 |
| Water viscosity (kg/nm/ns), $\eta$ | 10<sup>-21</sup> |
| Radius of protein (Å), $r_p$ | 13.5 |
| Radius of crowder C<sub>1</sub> (Å), $r_{c1}$ | 20 |
| Radius of crowder C<sub>2</sub> (Å), $r_{c2}$ | 12 |
| Diffusion constant of protein ($\mu\text{m}^2/\text{s}$), $D_p$ | 162.8 |
| Diffusion constant of C<sub>1</sub> ($\mu\text{m}^2/\text{s}$), $D_1$ | 109.9 |
| Diffusion constant of C<sub>2</sub> ($\mu\text{m}^2/\text{s}$), $D_2$ | 183.1 |

<sup>†</sup> Diffusion constants are calculated by using the hydrodynamic radii<sup>57</sup> in the Stokes-Einstein relation with viscosity of water $\eta = 1$ cP and $T = 300$ K.

Apart from attractive interactions, the macromolecular crowders inside the cytoplasm can also have repulsive interactions with the reacting proteins. Many polymeric crowders interact inertly and the effects of this kind of inert/repulsive crowders on protein association can be analytically obtained from Scaled Particle Theory(SPT). ^10,16,76–79^ To compare with SPT, protein crowder-interaction is modeled using a harmonic repulsive (HR) potential *V*_*HR*_(*r*) of the form,

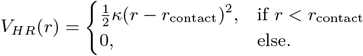

Here *κ* denotes the force constant, *r* is the inter-particle distance and *r*_contact_ is the corresponding distance between particle centers at contact.

In binary mixture of two crowders C_1_ and C_2_ with different protein-crowder interaction potentials are used (HR and LJ), in various mixing ratios (1:0, 0:1, 3:1 and 1:3) and at four crowder volume fractions (0.05, 0.15, 0.25 and 0.35) to systematically investigate the effect of various mixing environments on protein-protein association. Dimerization in mixing ratios 1:0 and 0:1 represent single crowder environments which were simulated to study the individual effects of the crowders.

During the simulation, the two proteins are in monomer state or in dimer state. Let 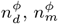 are the number of snapshots at equilibrium containing dimer and monomer species, respectively, at crowder volume fraction *ϕ*, and 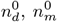 are the corresponding number of snapshots without crowders. Then the free energy difference for dimer formation can be directly evaluated as ^80,81^

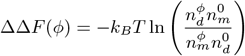

In ReaDDy, the GB1 dimerization happens with a microscopic rate proportional to experimental association rate when both the protein centers are within the reaction volume. The details of calculation of microscopic association and dissociation rates is provided in the Supporting Information. Each simulation at a particular volume fraction *ϕ* was run for 120 *µ*s with a time step of Δ*t* = 1 ps. ^82^ To calculate ΔΔ*F*, snapshots containing dimer or monomer protein species were saved every 100 steps. Error bars were estimated by running eight independent trajectories at each volume fraction and calculating the root mean square deviation from the average. In aggregate, 65.28 ms of simulation trajectory was generated.

## 3 Results

Inert repulsive crowders stabilize formation of dimer, as shown previously in both simulations and experiments. ^14,36,37,41^ However, we demonstrate below that attractive crowders can either stabilize or destabilize depending on a variety of factors including crowder volume fraction, crowder size and the strength of attraction. This has crucial implications in a binary crowder mixture.

### Stabilization or destabilization by attractive crowders

To investigate the effect of protein-crowder attraction strength on the GB1 protein dimer stability at a crowder volume fraction of 0.07 used in experiments, ^14^ we varied the well-depth *ϵ* of the LJ potential between the protein-crowder pair. In presence of large (20 Å) crowders at *ϕ* = 0.07, although there is no destabilization observed on average, the stabilization follows a non-linear trend (Fig. 2a *green triangles*). The stabilization decreases as the protein-crowder attraction becomes stronger up to a value of *ϵ* = 8 kJ/mol. With further increase in *ϵ*, the stabilization is increased. In presence of smaller (12 Å) attractive crowders, the destabilization effect is more pronounced at the same volume fraction *ϕ* = 0.07 (Fig. 2a *yellow squares*). The stabilization in this case initially decreases, with destabilization occurring between *ϵ* = 3-8 kJ/mol, and stabilize again for higher *ϵ*.

**Figure 2:**
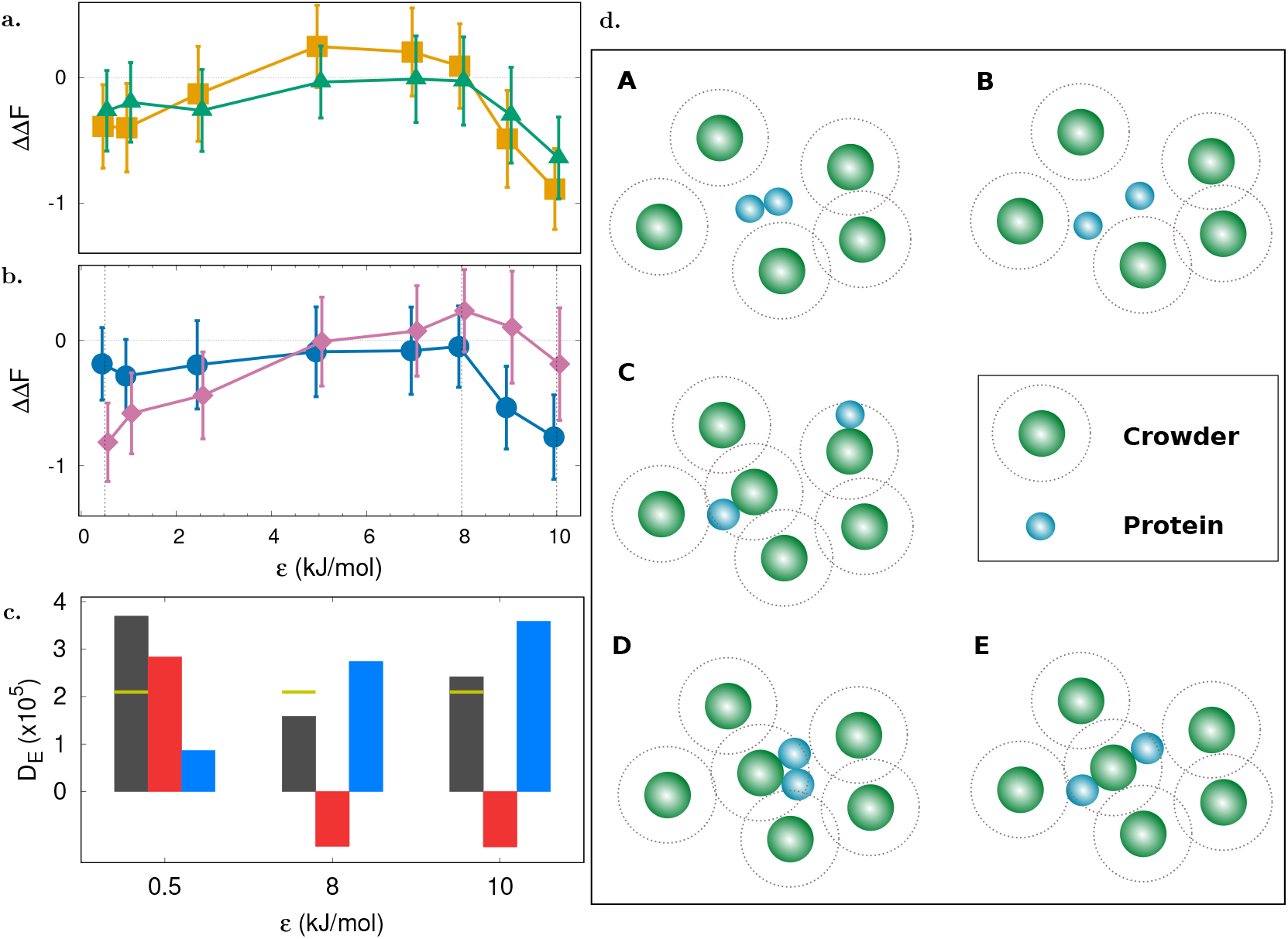
(a) ΔΔ*F* vs. *ϵ* in single crowder environment of 20 Å (*green triangles*) and 12 Å (*yellow squares*) crowders at *ϕ*=0.07. (b) ΔΔ*F* vs. *ϵ* in single 20 Å crowder at *ϕ* = 0.05 (*blue circles*) and 0.15 (*pink diamonds*). Vertical dotted lines show the values of *ϵ* studied for which ΔΔ*F* is maximum/minimum. (c) Total excess dimer frames (*D*_*ex*_) (*gray bars*), excess dimer frames simultaneously outside the crowder cutoff 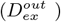 (*red bars*), and, excess dimer frames simultaneously inside the crowder cutoff 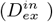 (*blue bars*) in presence of 20 Å crowders at different attraction strengths. The *yellow* line marks *D*_*ex*_ for *ϕ* = 0 case. (d) Configurations of proteins (*blue spheres*) and 20 Å crowders (*green spheres*) occurring during the simulations. The dotted line around the crowder indicates the cutoff. All energy units are in kJ/mol.

Crowder volume fraction also affects stabilization or destabilization at different *ϵ*. Fig. 2b compares the ΔΔ*F* versus *ϵ* for the larger crowder at two volume fractions, *ϕ* = 0.05 (Fig. 2b *blue circles*) and 0.15 (Fig. 2b *pink diamonds*). At the higher volume fraction destabilization begins at *ϵ* ~ 5 kJ/mol and the highest destabilization is observed at *ϵ* = 8 kJ/mol, after which the profle turns downward towards stabilization again.

To investigate the non-monotonic variation of ΔΔ*F* as a function *ϵ* in further detail let us consider the configurations of the GB1 protein in monomer and dimer states for crowder species (for example, the large 20 Å crowder) during the simulation (Fig. 2d). In presence of attractive crowders monomer or dimer species may form inside or outside the LJ cutoff of the crowder. Let, 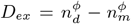 denote the difference in the number of snapshots having dimer and monomer species (or excess dimer species snapshots) at any distance from crowders, whereas 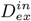 and 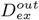 denote the same when dimers or monomers occur *simultaneously* inside or outside the LJ cutoff distance from any crowder, respectively. Thus, dimers and monomers forming *simultaneously* outside the cutoff distance from any crowder are represented by configurations (A-C), whereas they form *simultaneously* inside in configurations (D,E).

For elucidation of the ΔΔ*F* variation, consider the larger (20 Å) crowders at *ϕ* = 0.15 (Fig. 2b *pink diamonds*) at *ϵ* values 0.5, 8 and 10 kJ/mol, showing −ve, +ve (highest) and −ve ΔΔ*F*, respectively, while Fig. 2c shows the 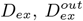 and 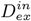 at the three *ϵ* values. In simulations, at low attraction strengths (*ϵ* = 0.5 kJ/mol), most of the monomer and dimer configurations occur outside the LJ cutoff distance so that the major contribution to ΔΔ*F* comes from 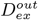 (entropic stabilization due to excluded volume). In Fig. 2d, configurations A and (B,C) depict corresponding dimer and monomer configurations, respectively. Stronger interaction (*ϵ* = 8 kJ/mol) between protein crowder species results in more monomers coming inside cutoff of different crowders (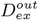 is −ve), as well as more dimers occurring inside the cutoff (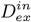 is +ve). After the monomers come within the cutoff of a particular crowder, they can come in contact to form the dimer (configuration D) or stay separate (configuration E). However, since in this case less dimers form than monomers compared to the dilute medium (no crowders), a net destabilization (ΔΔ*F* is +ve) is observed. In Fig. 2c, a higher (lower) *D*_*ex*_ (*gray bar*) than the reference (*yellow* line marks for *D*_*ex*_ at *ϕ* = 0) indicates overall stabilization (destabilization) of dimer. Here, it is to be noted that monomer and dimer configurations are in dynamic equilibrium during simulations, that is, they form and dissociate. A higher *ϵ* helps the monomers to stay within the crowder cutoff for a longer time, increasing the probability of dimer formation. With further increase in interaction strength (*ϵ* = 10 kJ/mol), although there is no change in the monomer or dimer configurations outside the cutoff, it becomes more likely that the monomers inside the cutoff come in contact with each other to form the dimer, showing a rise in 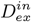 and an overall increase in *D*_*ex*_ that ensues stabilization.

### Dimerization in single crowders

Before investigating the mixed crowder environments, the individual crowder effects on the protein dimerization process is studied to verify the qualitative ΔΔ*F* versus *ϕ* trends (Fig. 3). For protein-crowder repulsive interaction, a harmonic repulsive potential having a force constant *κ* = 40 kJ/mol is used, whereas, for attractive interaction between the protein and the larger crowder, the LJ interaction strength is taken as *ϵ* = 8 kJ/mol, i.e., the strength at which maximum destabilization is observed as shown in Fig. 2b. A weak attraction strength of *ϵ* = 1 kJ/mol is used between the protein and smaller crowder to mimic a sticky environment.

**Figure 3:**
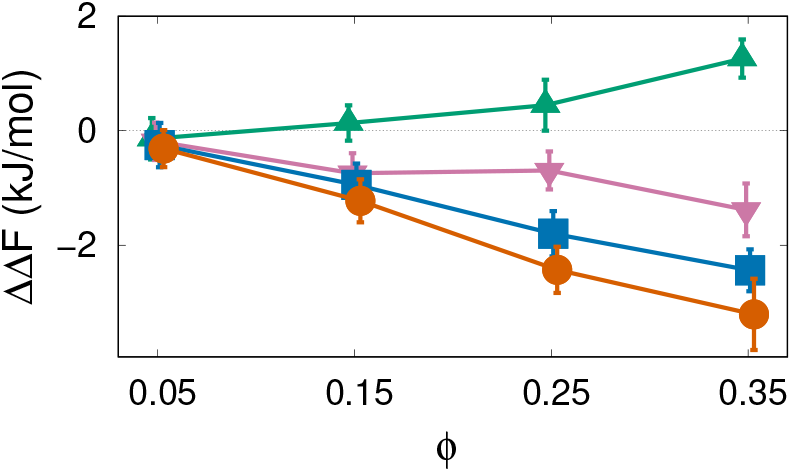
Change of free energy ΔΔ*F* for single crowder environments with various protein-crowder pairwise potentials: 20 Å crowder interacting with Lennard-Jones potential (*ϵ* = 8 kJ/mol) (*green up-triangles*) and harmonic repulsive potential (*κ* = 40 kJ/mol) (*blue squares*), 12 Å crowder interacting with Lennard-Jones potential (*ϵ* = 1 kJ/mol (*pink down-triangles*) and harmonic repulsive potential (*orange circles*).

In presence of large (20 Å) repulsive crowders, the stabilization of protein dimer increases with *ϕ* (Fig. 3 *blue squares*). The dimer configuration is entropically favored as it increases the available volume to the 20 Å crowders, in line with the prediction by the SPT. ^76,79^ A greater stabilization is observed in presence of small (12 Å) repulsive crowders (Fig. 3 *orange circles*) compared to that of the larger crowders (*blue squares*), as reported previously. ^56,83,84^ The comparison of ΔΔ*F* for repulsive crowder environments with SPT calculations is given in Supporting Information (Fig. S5).

Attractive interactions between protein and 20 Å crowders destabilize the protein dimer formation at higher *ϕ* (Fig. 3 *green up-triangles*). A weakly attractive protein-12 Å crowder interaction shows reduced stabilization than an environment of repulsive protein-12 Å crowder interaction (Fig. 3 *pink down-triangles*). With a higher protein-12 Å attraction strength of *ϵ* = 5 kJ/mol, destabilization increases with *ϕ* (see Supporting Information Fig. S9).

### Dimerization in a binary mixture

*In vivo* proteinprotein association occurs in a heterogeneous surrounding, having diverse interactions and conditions governing the overall behavior. A binary mixture model, although not replicating the complex cellular environment, provides a systematic step towards understanding the influence of these different conditions. As mentioned before, here the dimerization free energy is investigated in a mixture of two crowders: One (R_*h*_ ~ 20 Å) bigger and the other (R_*h*_ ~ 12 Å) smaller than the protein monomers (R_*h*_ 13. ~ 5 Å). Crowders can have either repulsive (HR) or attractive (LJ) interaction with the protein monomers. All crowder-crowder interactions are repulsive in all cases. The force constant of HR potential is *κ* = 40 kJ/mol for both crowder species. For attractive protein-crowder interaction, the well-depths of LJ potentials are chosen to be *ϵ* = 8 kJ/mol and *ϵ* = 1 or 5 kJ/mol for 20 Å and 12 Å crowder species, respectively. Thus, four different combinations of interactions between protein monomers and crowders are studied at two mixing ratios as detailed in Table. 2. For each pair of interactions, at specific total volume fraction (*ϕ*) of the crowders, the mixing ratio of 3:1 (1:3) denotes that the volume fractions of the 20 Å and 12 Å crowders are 0.75*ϕ* (0.25*ϕ*) and 0.25*ϕ* (0.75*ϕ*), respectively.

**Table 2:** Interaction potential between protein-20 Å crowder (C_1_) and protein-12 Å crowder (C_2_) respectively.

| Cases | Protein-crowder($C_1$) interaction potential | Protein-crowder($C_2$) interaction potential |
| --- | --- | --- |
| $\kappa - \kappa$ | HR ($\kappa = 40$ kJ/mol) | HR ($\kappa = 40$ kJ/mol) |
| $\kappa - \epsilon$ | HR ($\kappa = 40$ kJ/mol) | LJ ($\epsilon = 1$ kJ/mol) |
| | HR ($\kappa = 40$ kJ/mol) | LJ ($\epsilon = 5$ kJ/mol) |
| $\epsilon - \kappa$ | LJ ($\epsilon = 8$ kJ/mol) | HR ($\kappa = 40$ kJ/mol) |
| $\epsilon - \epsilon$ | LJ ($\epsilon = 8$ kJ/mol) | LJ ($\epsilon = 1$ kJ/mol) |
| | LJ ($\epsilon = 8$ kJ/mol) | LJ ($\epsilon = 5$ kJ/mol) |

### Case I: HR-HR (*κ* - *κ*) interaction between protein and crowders

In the binary crowder environment, where both 20 Å and 12 Å crowders have repulsive interactions with GB1 proteins (Fig. 4), the 3:1 mixing composition (Fig. 4 *olive up-triangles*) nearly follows the single 20 Å crowder trend (Fig. 4 *blue squares*). Interestingly, for 1:3 mixing composition (Fig. 4 *red down-triangles*), that is, when the 12 Å crowder is the dominant crowding species, the stabilization greatly decreases compared to single 12 Å crowder (Fig. 4 *orange circles*) case. This can be understood in terms of the reduction in excluded volume in presence of crowders of various sizes. For single crowders (say, the 12 Å crowder), the volume excluded when the protein-protein pair is in dimer configuration is less (or available volume is more) compared to the monomer configuration, and it is entropically favorable to bring the protein monomers together to form the dimer state.

**Figure 4:**
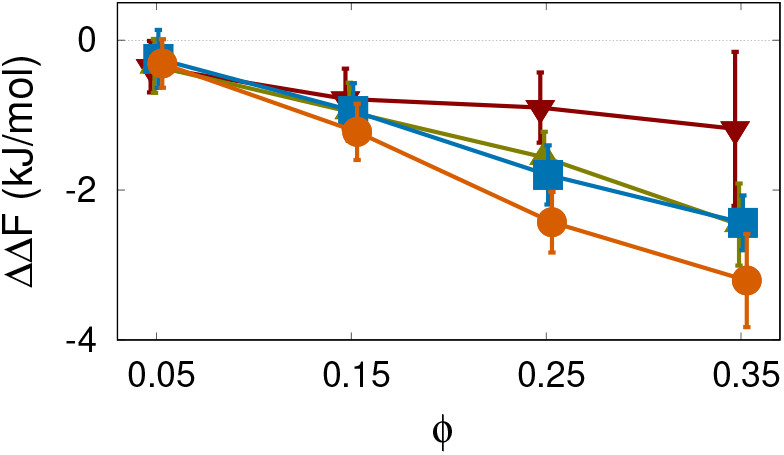
Dimerization in presence of mixed crowders interacting via repulsive potentials. Free energy changes for single crowders, 20 Å (*blue squares*) and 12 Å (*orange circles*), and in mixing ratio 3:1 (*olive up-triangles*) and 1:3 (*red down-triangles*).

Similarly, when the 12 Å crowder is the dominant species in a mixture of crowders of two different sizes (12 Å and 20 Å), it will try to bring all other possible pairs of species (20 Å-20 Å crowder-crowder, 13.5 Å-20 Å protein-crowder and 13.5 Å-13.5 Å protein-protein) together to minimize the excluded volume. Considering the particles as hard spheres, the reduction in the volume excluded to 12 Å crowder, when the other species are in contact and stay separate is in the order (highest to lowest) of C_1_-C_1_, protein-C_1_, and proteinprotein (see Supporting Information Table. S2). There is no direct effect on ΔΔ*F* when the larger crowders C_1_-C_1_ come in contact. But when the protein-20 Å pair is favored over the protein-protein pair, the stability of the protein dimer state is reduced compared to the single crowders (Fig. 4). Still, there is stability of the protein dimer state as the monomers forming the dimer have a reaction, in analogy of the specific interaction between them, as compared to protein-20 Å pair interacting only via repulsive potential. The exact number of these configurations are shown in the Supporting Information Fig. S8.

### Case II: HR-LJ (*κ* - *ϵ*) interaction between protein and crowders

We next consider an environment in which the protein-smaller (12 Å) crowders have low attractive strength, while the protein-larger crowder (20 Å) interaction is still repulsive. Fig. 5a shows the overall change in free energy which also includes ΔΔ*F* for single crowder species for reference. Fig. 5b and c shows the enthalpic and entropic contributions to ΔΔ*F*, respectively. Fig. 5d depicts major monomer (F) and dimer (G) configurations. Fig. 5e and f plots the number of excess dimer configurations inside and outside of the cutoff distance of the smaller crowder at 3:1 and 1:3 crowder composition, respectively, providing an outlook of their contribution on the enthalpic and entropic stabilization.

**Figure 5:**
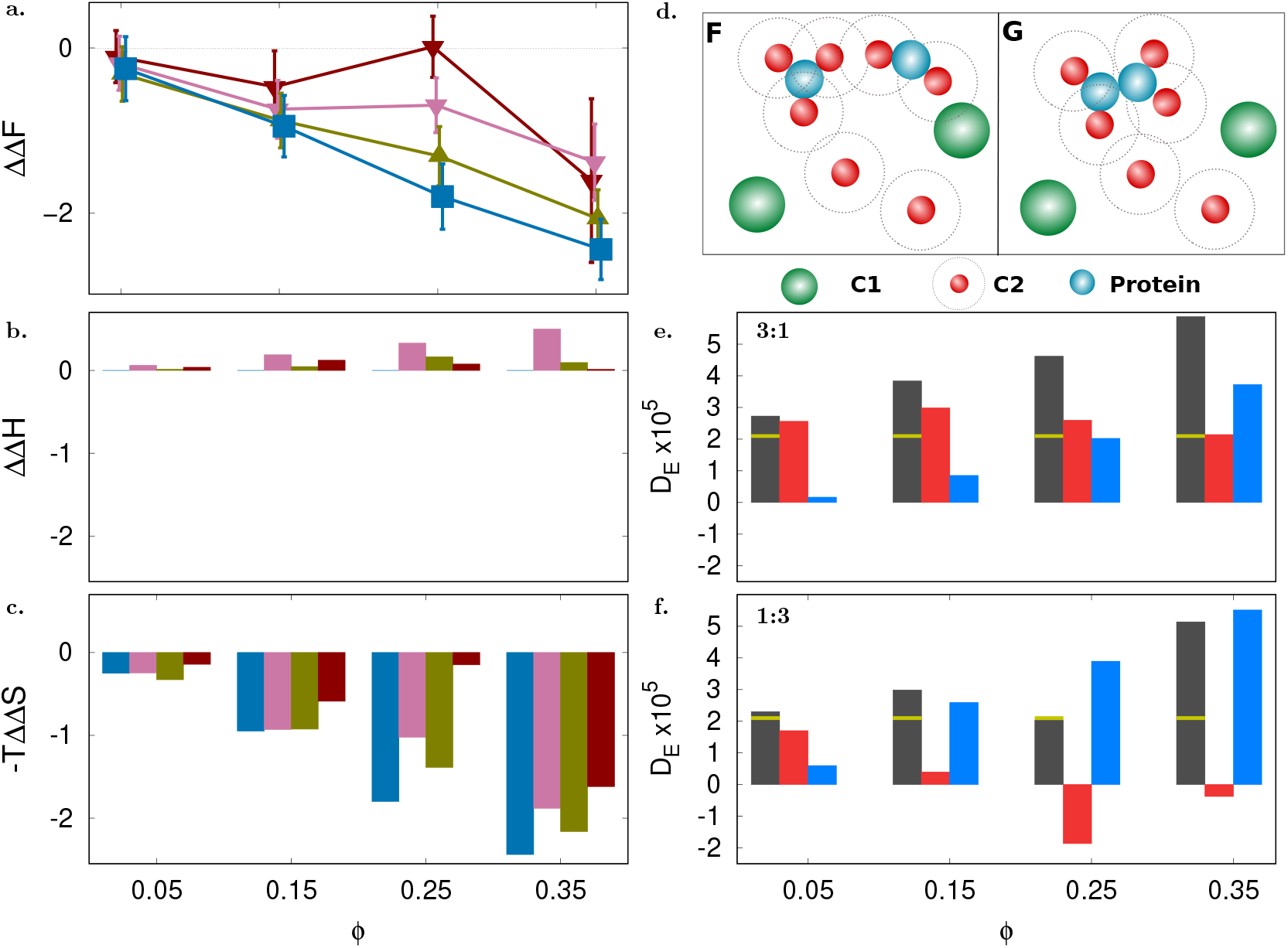
Dimerization in presence of mixed HR-LJ interactions. (a) Change in free energy ΔΔ*F* in presence of a mixed environment of 20 and 12 Å crowders at mixing ratio 3:1 (*olive up-triangles*) and 1:3 (*red down-triangles*) interacting via HR and LJ potentials, respectively. ΔΔ*F* for single crowder environments with protein-20 Å HR potential (*blue squares*), protein-12 Å LJ (*ϵ* = 1 kJ/mol) potential (*pink down-triangles*) are shown for reference. (b) Bars showing change in enthalpy ΔΔ*H* at each volume fraction *ϕ* in the order of single 20 Å, 12 Å crowders and mixed crowders in 3:1 and 1:3 ratios (bars follow the line colors defined in (a)). (c) Bars showing change in entropy −*T* ΔΔ*S* at each volume fraction (as in (b)). (d) Major configurations contributing to the monomer and dimer frames. (e) Bars indicating excess dimers for 3:1 mixing ratio shown in the order of 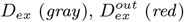, and 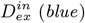. The *yellow* line marks *D*_*ex*_ for *ϕ* = 0 case. (f) Same quantities as (e) for 1:3 mixing ratio. All energy units are in kJ/mol.

For 3:1 mixing ratio, dimer formation is stabilized for all volume fractions (Fig. 5a *olive up-triangles*). At this composition, it is entropically favored to bring protein monomers together by 20 Å repulsive crowders. Also, two or three small sticky crowders can transiently interact with the protein which acts as an unit (Fig. 5d). With increase in *ϕ*, it is entropically favored to bring the protein monomers or transient protein-12 Å crowder units together (Fig. 5c *olive bar*) which can lead to formation of dimer. The corresponding monomer and dimer configurations are shown in Fig. 5d. This can also be seen with the increase in 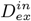 with *ϕ* (Fig. 5e *blue bars*) showing increase in dimer configurations within the cutoff of 12 Å crowders and reduction in the dimer configuration forming outside the cutoff (Fig. 5e *red bars*). Hence, the overall stabilization is entropic at 3:1 mixing ratio. The reduced stabilization compared to single 20 Å crowder environment (Fig. 5a *blue squares*) can arise due to entropic stabilization in bringing protein-12Å transient unit and 20 Å crowder together.

At 1:3 composition, the 12 Å crowders are weakly interacting but occupies more volume than the larger crowder. Accordingly, ΔΔ*F* (Fig. 5a *red down-triangles*) has a small enthalpic but large entropic component (Fig. 5c *red bars*). With more 12 Å crowders, it is entropically more favored to bring the protein-12 Å transient units together. This leads to an increase in 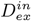 inside the cutoff of 12 Å crowders and decrease in 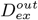 (*blue* and *red* bars in Fig. 5f). At lower volume fractions, the entropic effect can stabilize both protein-12 Å transient unit and 20 Å crowder together or two protein-12 Å transient units together. At the highest *ϕ*, entropic stabilization for the protein-12 Å crowder units, is highest. In case of stronger attraction (*ϵ* = 5 kJ/mol) between protein and the smaller 12 Å crowder, enthalpic contribution shifts the ΔΔ*F* profile upwards as shown in Supporting Information (Fig. S10).

### Case III: LJ-HR (*ϵ* - *κ*) interaction between protein and crowders

Consider a binary crowder mixture in which the larger 20 Å crowder interacts with the protein via a strong attractive potential (*ϵ* = 8 kJ/mol) whereas the smaller crowder interacts inertly via a repulsive potential. This crowding scenario is likely if, for example, the protein dimerization is observed in presence of large attractive protein (say, lysozyme) and inert polymer (say, ficoll or dextran of low molecular weight) crowder species. Fig. 6a-f describes similar quantities as in the previous case. As the large crowder is attractive, 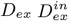, and 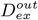 are calculated w.r.t cutoff of 20 Å crowder.

**Figure 6:**
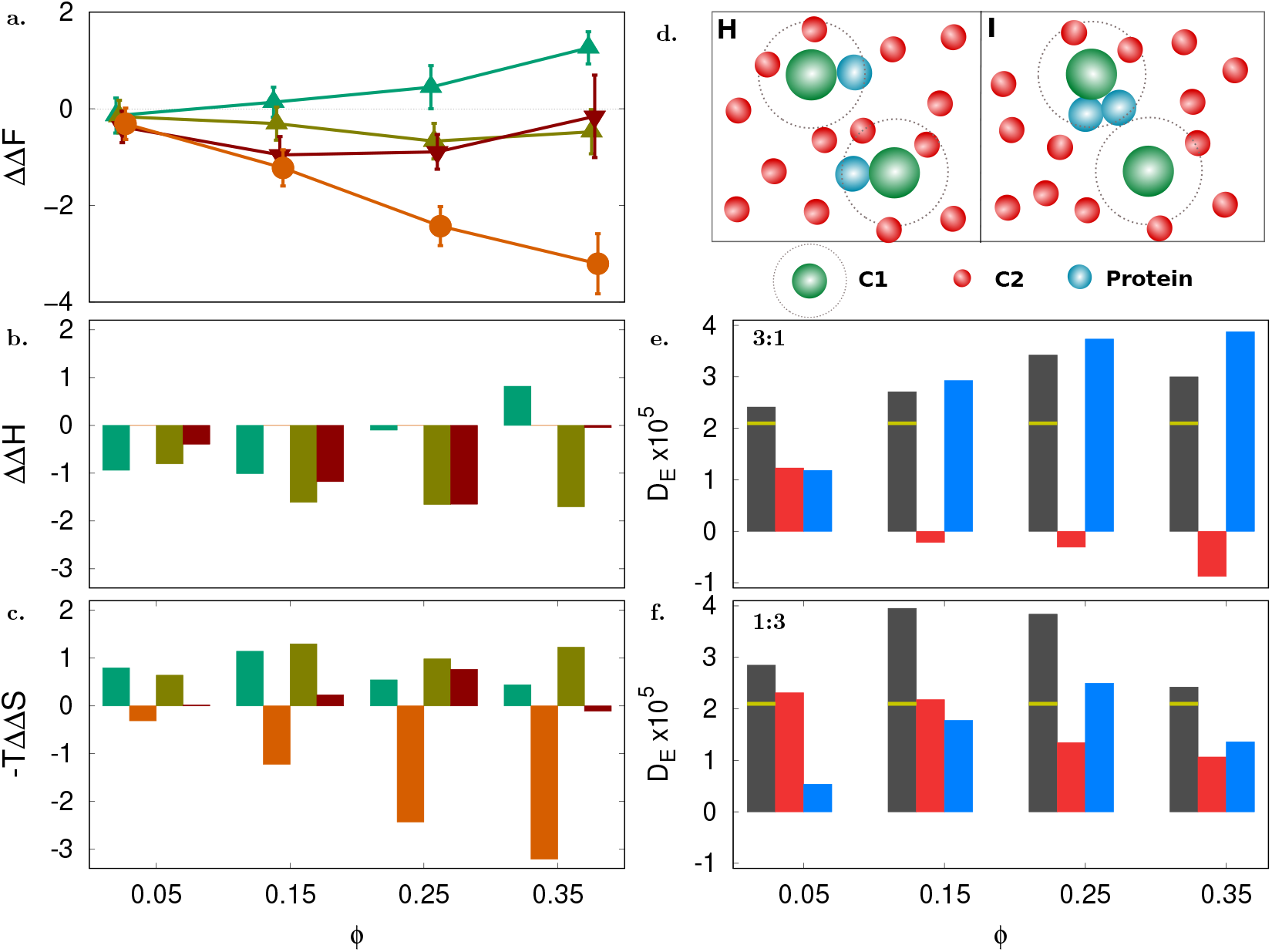
Dimerization in presence of mixed LJ-HR interactions. (a) Change in free energy ΔΔ*F* in presence of a mixed environment of 20 and 12 Å crowders at mixing ratio 3:1 (*olive up-triangles*) and 1:3 (*red down-triangles*) interacting via LJ and HR potentials, respectively. ΔΔ*F* for single crowder environments with protein-20 Å LJ (*ϵ* = 8 kJ/mol) potential (*green up-triangles*), protein-12 Å HR potential (*orange circles*) are shown for reference. (b) Bars showing change in enthalpy ΔΔ*H* at each volume fraction *ϕ* in the order of single 20 Å, 12 Å crowders and mixed crowders in 3:1 and 1:3 ratios (bars follow the line colors defined in (a)). (c) Bars showing change in entropy −*T* ΔΔ*S* at each volume fraction (as in (b)). (d) Major configurations contributing to the monomer and dimer frames. (e) Bars indicating excess dimers for 3:1 mixing ratio shown in the order of 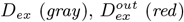, and 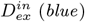. The *yellow* line marks *D*_*ex*_ for *ϕ* = 0 case. (f) Same quantities as (e) for 1:3 mixing ratio. All energy units are in kJ/mol.

For 3:1 mixing ratio, dimer formation is stabilized for all volume fractions (Fig. 6a *olive up-triangles*). In this composition, at low *ϕ*, it is enthalpically favored to form dimer (Fig. 6b *olive bars*) similar to single attractive 20 Å crowder (Fig. 6b *green bars*). Additionally, after the formation of protein dimer-20 Å crowder configuration I (Fig. 6d) it is entropically stabilized by the 12 Å repulsive crowders. This entropic stabilization of configuration I increases the enthalpic stabilization of dimers by 20 Å crowders even at higher volume fractions unlike in the case of single attractive 20 Å crowders. This can also be seen with the increase in 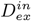 with *ϕ* (Fig. 6e *blue bars*) showing increase in dimer configurations within the cutoff of 20 Å crowders and reduction in the dimer configuration forming outside the cutoff (Fig. 6e *red bars*). Hence the overall enthalpic stabilization is sustained by entropy.

At 1:3 composition, the 12 Å repulsive crowders occupy more volume than the attractive large crowder. Accordingly, ΔΔ*F* (Fig. 6a *red down-triangles*) has less enthalpic stabilization contribution (Fig. 6b *red bars*). Apart from enthalpic stabilization of attractive 20 Å crowders, the small repulsive crowders can entropically stabilize protein dimer or protein-20 Å crowder together. As there are more 12 Å repulsive crowders in 1:3 composition, the 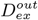 (Fig. 6f red bars) is more than that in 3:1 composition (Fig. 6e red bars) while 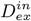 (Fig. 6f *blue bars*) is less compared to 3:1 composition (Fig. 6e *blue bars*).

### Case IV: LJ-LJ (*ϵ* - *ϵ*) interaction between protein and crowders

Finally, consider both crowders having attractive LJ interaction with the protein. The protein-20 Å crowder has attraction strength of 8 kJ/mol, while the 12 Å crowder has a strength of 1 kJ/mol, mimicking small cellular components with ‘sticky’ interactions. This crowding scenario explores dimerization in an environment of competitive non-specific binders. Again, Fig. 7a-f describes similar quantities as in the previous case.

**Figure 7:**
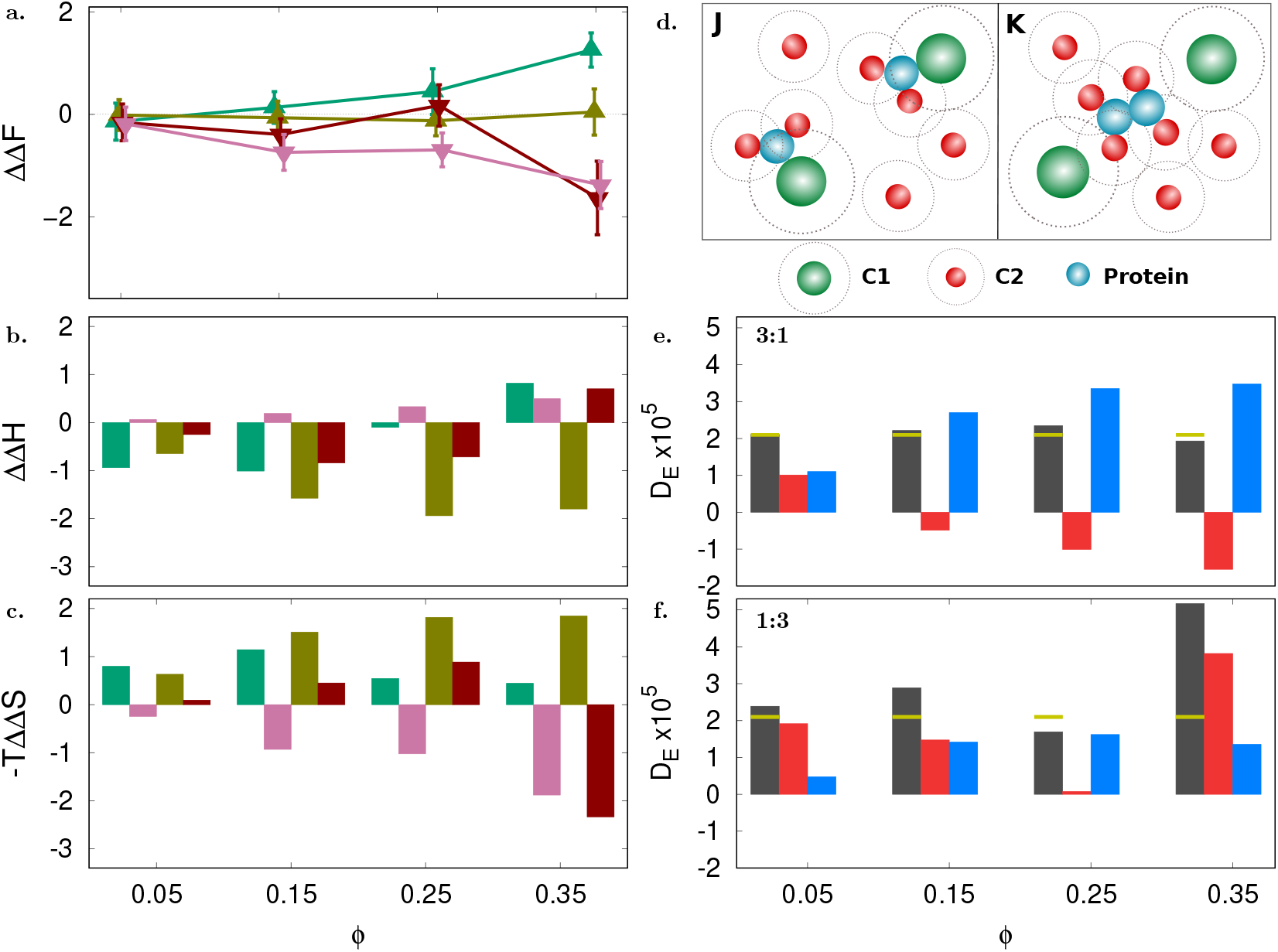
Dimerization in presence of mixed LJ-LJ interactions. (a) Change in free energy ΔΔ*F* in presence of a mixed environment of 20 and 12 Å crowders at mixing ratio 3:1 (*olive up-triangles*) and 1:3 (*red down-triangles*) interacting via LJ and LJ potentials, respectively. ΔΔ*F* for single crowder environments with protein-20 Å LJ (*ϵ* = 8 kJ/mol) potential (*green up-triangles*), protein-12 Å LJ (*ϵ* = 1 kJ/mol) potential (*pink down-triangles*) are shown for reference. (b) Bars showing change in enthalpy ΔΔ*H* at each volume fraction *ϕ* in the order of single 20 Å, 12 Å crowders and mixed crowders in 3:1 and 1:3 ratios (bars follow the line colors defined in (a)). (c) Bars showing change in entropy −*T* ΔΔ*S* at each volume fraction (as in (b)). (d) Major configurations contributing to the monomer and dimer frames. (e) Bars indicating excess dimers for 3:1 mixing ratio shown in the order of 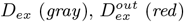, and 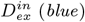. The *yellow* line marks *D*_*ex*_ for *ϕ* = 0 case. (f) Same quantities as (e) for 1:3 mixing ratio. All energy units are in kJ/mol.

For 3:1 mixing ratio, dimer formation stability is nearly zero (Fig. 7a *olive up-triangles*). At this composition, with increase in *ϕ*, it is enthalpically favored (Fig. 7b *olive bars*) for dimer to form inside the cutoff of 20 Å crowder. This can be seen with increase in 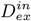 with *ϕ* (Fig. 7e *blue bars*). It is also entropically favored to bring the protein-12 Å crowder transient unit and 20 Å crowder together, resulting in entropic destabilization (Fig. 7c *olive bars*) of protein dimer and reduction of excess dimer frames 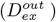 outside the cut-off of 20 Å crowders (Fig. 7e *red bars*). The overall effects are canceled out giving similar results as in an environment without any crowder.

At 1:3 composition, the weakly interacting 12 Å crowders occupy more volume than 20 Å attractive crowders. Accordingly, ΔΔ*F* (Fig. 7a *red down-triangles*) has less enthalpic stabilization (Fig. 7b *red bars*) due to 20 Å crowder than in 3:1 composition. This can be seen by increase in 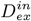 (Fig. 7f *blue bars*) which is less in magnitude compared to 3:1 composition. More entropic stabilization is observed by bringing two protein-12 Å transient units together. At lower volume fractions, there is competition between bringing protein-12 Å transient unit and 20 Å crowder together or bringing two protein-12 Å transient units together. This decreases the 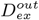 (Fig. 7f *red bars*) up to *ϕ* = 0.25. At the highest volume fraction, the entropic stabilization effect is highest due to large number of small crowders which stabilizes the biggest units, i.e protein-12 Å transient units together resulting in stabilization.

In case of stronger attraction (*ϵ* = 5 kJ/mol) between protein and the smaller 12 Å crowder, enthalpic contribution shifts the ΔΔ*F* profile upwards as shown in Supporting Information (Fig. S11).

## 4 Discussion

A crowded macromolecular environment is like a multicomponent soup of biological macromolecules. In addition to understanding the role of individual components of the soup a key interest in the field has been to find out if all the components together behave in a different manner and explain the observed differences. In this regard, many studies, both experimental and theoretical, indicate that crowding effects are non-additive, ^38,85,86^ although contrasting evidences have also been reported. ^54^ In this work we provide a detailed description of free energy changes under different crowding conditions. By systematic investigation of various binary crowding scenarios it is shown that the overall effect emerges due to complex balance of entropic and enthalpic contributions, which cannot be understood by considering the property of one crowder component in isolation.

Soft attractive interactions are known to destabilize protein-protein association. The destabilizing effect is seen for EIN-HPr binding in the presence of attractive BSA crowders as well as for GB1 dimerization in presence of lysozyme crowders. ^14^ Although our preliminary model of GB1 dimerization in presence of lysozyme crowders did not capture the destabilizing effect, by choosing a narrower modified form of LJ potential in this work, we recover the destabilizing influence of attractive crowders (Fig. 2a). In experiments, the ΔΔ*F* for side-by-side GB1 dimer formation is 0.76 ± 0.21 kJ/mol at *ϕ* = 0.07, whereas in this work we find ΔΔ*F* = −0.02 ± 0.38 kJ/mol at the same value of *ϕ* with *ϵ* = 8 kJ/mol. Destabilization is enhanced at higher volume fractions (ΔΔ*F* = 0.13 ± 0.31 kJ/mol at *ϕ* = 0.15) showing that the model can capture qualitatively same behavior as in experiments.

The primary goal of this study has been to investigate the influence of mixed crowder environments. Although mixed crowding has been shown to influence protein folding, ^38^ macromolecular domain movement, ^85^ enzyme activity and dynamics ^86^ and protein fibrillation, ^70^ few studies investigated protein association in a mixed crowded environment. ^56,57^ Previous all-atom cytoplasmic model of M. genitalium showed that stability of protein association is reduced in presence of a heterogeneous crowding environment compared to homogeneous crowding, similar to our observation for a binary mixture of two repulsive crowders (Fig. 4). It has also been shown that *in vivo* (cell lysate) and in lysis buffer protein-RNA binding is destabilized which otherwise was stabilizing in PEG crowders. ^37^ The reduction in binding affinity is attributed to increased dissociation rate due to the sticky components of the cellular matrix. In our simulations the sticky environment is mimicked by the smaller 12 Å crowder with low attraction strength. At large volume fraction (1:3) this environment induces reduced stabilization or destabilization depending on the strength of sticky interactions (Fig. 7 and SI Fig. S11). Further verification of results presented here require experiments in different mixed crowder conditions.

The non-monotonic effect of ΔΔ*F* .vs. *ϕ* in mixed crowder solutions, especially when the attractive smaller crowder is in higher volume fraction (Fig. 7 and SI Fig. S11), can arise due to the entropic effect stabilizing the protein-12 Å transient units (e.g Fig. 7d configuration K). In general, upto a certain volume fraction, here 0.25, the individual transient units are relatively stable resulting in reduced stabilization. At the highest volume fraction, the entropic effect overcomes this and stabilizes the protein dimer resulting in enhanced stabilization. Non-monotonic stabilization of a bridged state of cylindrical rods by small crowders (w.r.t without crowder medium) has been observed previously, ^17^ where the bridged state was stabilized up to a certain volume fraction after which the stabilization was reduced. To note that, in our study, the ΔΔ*F* of protein dimerization in crowder, as opposed to the bridged state in the previous study, is calculated w.r.t without crowder medium.

In summary, this work demonstrates that crowder mixing conditions regulate stability of protein dimerization. We also found that the emergent behavior of a mixed macromolecular environment can be deciphered through understanding of the entropic and enthalpic influences of crowders on other species in the surrounding, providing a qualitative idea on the trend of stability of protein-protein association in such environments. In cell, smaller crowders are likely to be present in large proportions. In our simulations it is observed that the smaller crowders have the dominant effect on ΔΔ*F* when present in larger volume fraction. In future, we would like to incorporate shape and site specific crowder interaction in our model. Also, it would be interesting to compare our observations against different experimental mixing conditions in future.

## Associated Content

## Supporting Information

Fitting GB1-lysozyme PMF to Lennard-Jones type potential. ReaDDy simulation parameters. Scaled Particle Theory (SPT). Excluded volume calculation. ΔΔ*H* Calculation. Single crowder environment with strong protein-12 Å LJ interaction (*ϵ* = 5 kJ/mol). Mixed crowder simulation with strongly attractive small (12 Å) crowders (*ϵ* = 5 kJ/mol)

## Acknowledgment

The authors gratefully acknowledge access to HPC facilities of NIT Rourkela, IIT Kanpur, IIT Kharagpur and Bioinformatics Resources and Applications Facility (BRAF), C-DAC, Pune for running the simulations. This work has been supported by the Department of Biotechnology (Grant No. BT/PR48048/BID/7/1026/2023), Government of India.

