## Supporting Information for "Emergent Effect of Mixed Crowders on Protein Dimerization"

### for

#### 1 Fitting GB1-lysozyme PMF to Lennard-Jones type potential

In order to determine the interaction potential between the GB1 protein and lysozyme crowders, a system with one GB1 protein and one lysozyme protein was prepared initially by coarse-graining the all-atom model using the MARTINI 3 force field.[1] While coarse-graining, harmonic bonds with a force constant of 500 kJ/mol/nm$^2$ were imposed on backbone of both GB1 and lysozyme, to ensure that their conformations remain close to their respective native state. The elastic network models of both GB1 and lysozyme were then introduced into a simulation box of dimension 90 x 90 x 90 Å$^3$ by using the PACKMOL tool.[2] With the GROMACS software,[3] the prepared system was solvated with the explicit MARTINI water. After neutralizing the net charge of the system with the addition of appropriate number of ions, a steepest descent energy minimization run was performed on the system. The energy minimization was followed by two NVT equilibrations, one with the positions of the atoms being restrained to their initial coordinates and another without any position restraints. Both the NVT equilibrations and the following NPT equilibration was executed for 50 ns. After the NPT equilibration, a production run of 500 ns was performed. This 500 ns trajectory was then analyzed for determining the variation of the distance between the center of geometry of GB1 and lysozyme using the "gmx pairdist" utility of the GROMACS software. From the probability distribution of center of geometry (CoG) distance between GB1 and lysozyme in the simulation, potential of mean force (PMF) was calculated by,

$$PMF(r) = -k_B \mathrm{T} \ln \left( \frac{p(r)}{p_{max}} \right)$$

where $p(r)$ = probability of CoG distance to be r and $p_{max}$ is the maximum probability.

Next, the shifted Martini PMF is fitted to the general Lennard-Jones potential with exponent m-n of the form,

$$V_{LJ}(r) = \frac{-\epsilon}{\left[ \left( \frac{\sigma}{r_m} \right)^m - \left( \frac{\sigma}{r_m} \right)^n \right]} \left[ \left( \frac{\sigma}{r} \right)^m - \left( \frac{\sigma}{r} \right)^n \right]$$

where $r_m$ is at contact distance of 33.5 Å, as shown in Fig. S1. We found m=48, n=28 exponents to be best suited residual sum of squares fitting parameters for the PMF.

The modified LJ (Kim-Mittal) potential[4] is given by,

$$V_{KH}(r) = 4 \left[ \epsilon_r \left( \frac{\sigma_r}{r - \sigma + \sigma_r} \right)^{12} - \epsilon_a \left( \frac{\sigma_r}{r - \sigma + \sigma_r} \right)^6 \right]$$

Here, the repulsive ($\epsilon_r$) and attractive ($\epsilon_a$) strengths control the narrowness of the LJ potential. This KM LJ potential can also be fitted (Fig. S2a) to the Martini PMF with $\epsilon_r = \epsilon_a = 8$ kJ/mol, which also minimizes the residual sum of squares (Fig. S2b). The interaction range $\sigma_r$ is set to 6 Å.

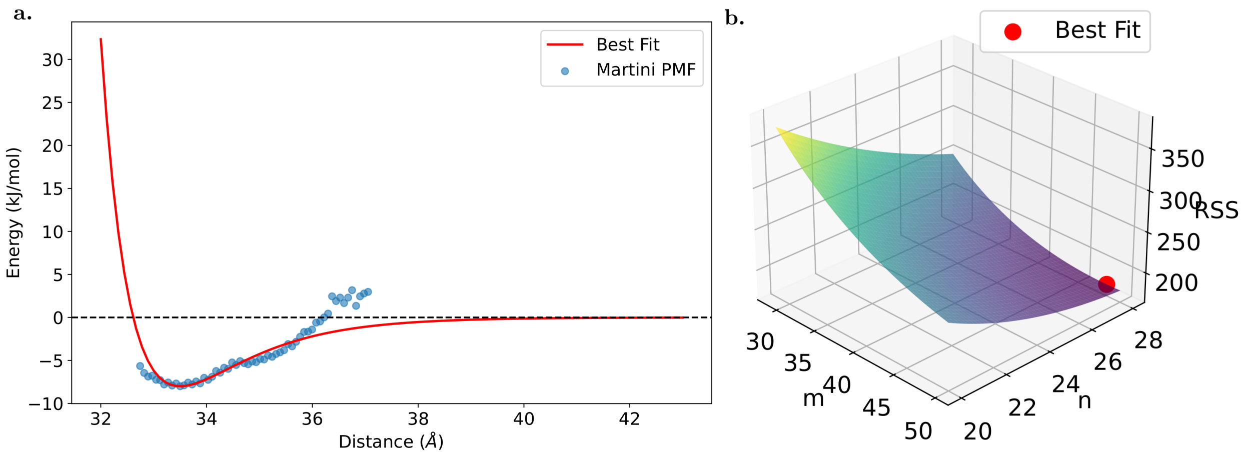


Figure S1: (a) Martini PMF and the fitted 48-28 LJ potential (red trace). (b) Residual sum of squares (RSS) for fit of LJ-type potential with exponents m-n to Martini PMF.

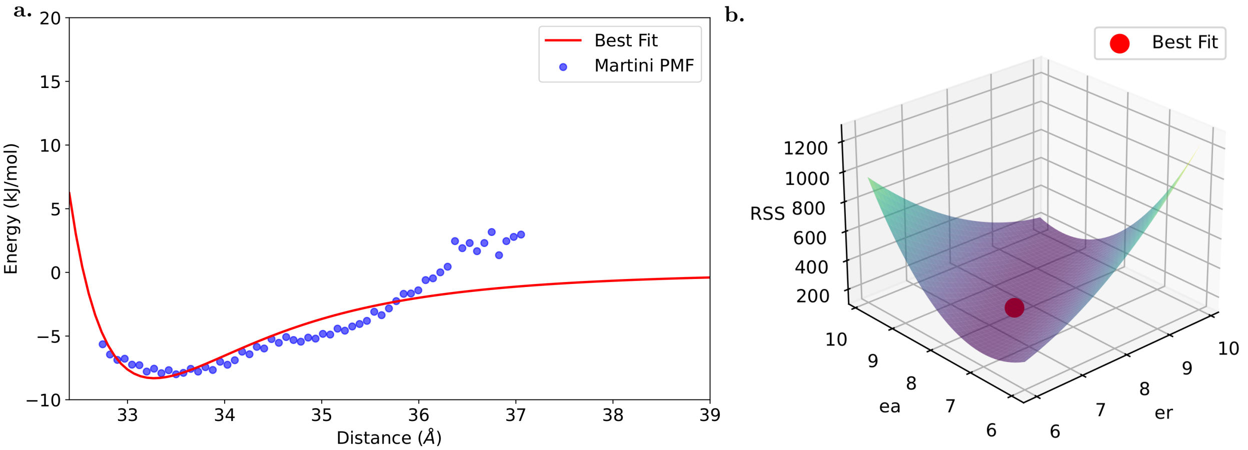


Figure S2: (a) Martini PMF and the fitted KM LJ potential. (b) Residual sum of squares for fit of KM potential to Martini PMF.

Fig. S3 shows a comparison between the fitted 48-28 LJ potential and the KM model.

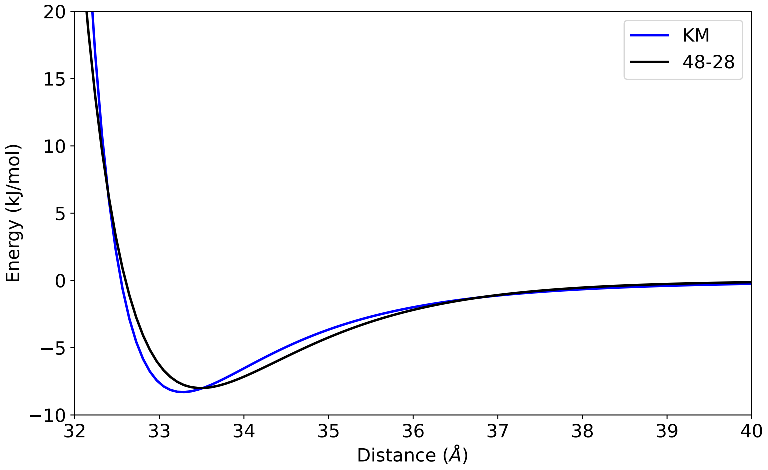


Figure S3: Comparison of 48-28 LJ-type potential with the KM LJ potential.

As the KM potential form matches closely, we use this potential to obtain m-n parameters for the protein-small crowder attractive interaction. A 40-20 exponent LJ potential matches closely with the corresponding KM potential for the protein-12 Å crowder interaction as shown in Fig. S4.

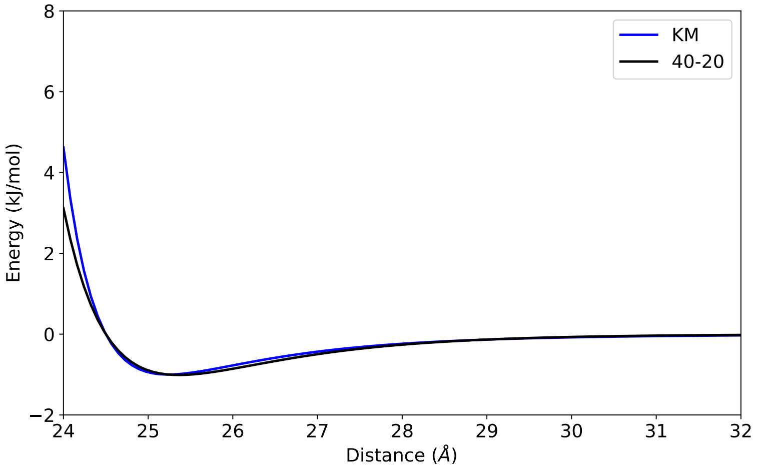


Figure S4: Comparison of 40-20 LJ-type potential with the KM LJ potential for 12 Å crowder.

As $r_m$ is at the contact distance of protein and crowder and the $\sigma$ at which the potential is zero becomes,

$$\sigma = \frac{r_m}{\left(\frac{m}{n}\right)^{\left(\frac{1}{m-n}\right)}}$$

Table S1: LJ-type model parameters for crowders of various sizes.

| Parameter | 20 Å | 12 Å | Unit |
|---|---|---|---|
| m | 48 | 40 | - |
| n | 28 | 20 | - |
| $r_{cutoff}$ | 1.4 $\sigma$ | 1.4 $\sigma$ | Å |
| $\epsilon$ | 8 | 1 or 5 | kJ/mol |
| $\kappa$ | 40 | 40 | kJ/mol |

#### 2 ReaDDy simulation parameters

**Microscopic association and dissociation rates($\lambda_{\text{on}}$ and $k_{\text{off}}$)**

In ReaDDy, GB1 proteins can diffuse and take part in reactions. The reaction events are discrete i.e. for each time step $\tau$, if the proteins are within a reaction radius, they can form the product with probability,[5,6]

$$p(\lambda;\tau) = 1 - e^{-\lambda_{\text{on}}\tau}$$

Here $\tau$ is the time step and $\lambda_{\text{on}}$ is the microscopic association rate, given by,

$$\lambda_{\text{on}} = \frac{k_{\text{on}}}{V_{\text{eff}}}$$

This equation relates the microscopic association rate with the macroscopic association rate $k_{\text{on}}$. When the two GB1 proteins are inside the reaction volume $V_{\text{eff}}$ given by,

$$V_{\text{eff}} = \int_0^R \exp(-\beta U) 4\pi r^2 \, dr$$

where R the reaction radius, they can associate with the microscopic association rate $\lambda_{\text{on}}$.

In this case, there is no interaction(U = 0) between the protein monomers except that they can associate with rate $\lambda_{\text{on}}$ forming the product if they come within the effective reaction volume given by,

$$V_{\text{eff}} = \frac{4}{3}\pi R^3$$

where, $R = R_p + R_p = 13.5 + 13.5 = 27$ Å is the contact distance between two GB1 monomers, Now the microscopic association rate becomes,

$$\lambda_{\text{on}} = \frac{k_{\text{on}}}{\frac{4}{3}\pi R^3}$$

The equilibrium dissociation constant $K_{\text{D}}$ is given by,

$$K_{\text{D}} = \frac{k_{\text{off}}}{k_{\text{on}}}$$

where, $k_{\text{off}}$ is the microscopic dissociation rate. The $K_D$ for GB1 dimerization is 59 $\mu$M.[7]

The GB1 dimer formed may undergo dissociation with the rate $k_{\text{off}}$.

We get the microscopic association rate as,

$$k_{\text{on}} = \frac{k_{\text{off}}}{K_{\text{D}}}$$

For $k_{\text{off}}$=2200 s$^{-1}$ we get $k_{\text{on}} = 37.3\ \mu\text{M}^{-1}s^{-1}$.

After rescaling the rates to observe many association and dissociation events in the simulation trajectory, we choose the microscopic association rate, $\lambda_{\text{on}} = 7.5\ ns^{-1}$ and microscopic dissociation rate, $\lambda_{\text{off}} = 2.2 \times 10^{-2}$ ns$^{-1}$.

**Timestep**

The timestep $\tau$ is obtained as[8] $\tau = 10^{-4}\tau_{\mathrm{d}}$, where $\tau_{\mathrm{d}} = \frac{R^2}{\mathrm{D}}$ is the time to traverse the reaction volume. The time step is taken sufficiently small for it to be acceptable for steep interaction potential of Lennard-Jones and is less than the inverse reaction rate.

**Calculation of $\Delta\Delta F$ from dimer and monomer state probabilities**

The equilibrium constant of association ($K_a$) for a finite system ($N_A = 2$) of particles, as obtained from a recent study $K_a = \frac{f^{A_2}}{2(1-f^{A_2})} V c^{\varphi}$,[9] gives the same expression for $\Delta\Delta F$ as taking the ratio of dimer and monomer fractions between crowder and without crowder medium the other factors drops out.

Let $x = \frac{p_0^0}{p_0^1}$ be the ratio of monomer and dimer probabilities in the absence of crowders and $y = \frac{p_1^\phi}{p_0^\phi}$ be the ratio of dimer and monomer probabilities in the presence of crowder volume fraction $\phi$. Then

$$\begin{aligned}
\Delta\Delta F &= \Delta F^{\phi} - \Delta F^{0} \\
&= -k_{\mathrm{B}} T \ln \left( \frac{p_1^\phi}{p_0^\phi} \frac{p_0^0}{p_1^0} \right) \\
&= -k_{\mathrm{B}} T \ln(xy)
\end{aligned}$$

and the error is calculated as

$$\sigma_{\Delta\Delta F} = \left( \frac{{\sigma_x}^2}{x^2} + \frac{{\sigma_y}^2}{y^2} \right)^{\frac{1}{2}}$$

#### 3 Scaled Particle Theory (SPT)

The free energy change in placing a spherocylinder with cylindrical length $2L_cR_p$ and hemispheres on each side with radius $R_p = R_c r_c$ with $r_c$ being the crowder radius is[10–12]

$$\frac{\Delta F^{\phi}}{RT} = -ln(1-\phi) + \alpha_1\xi + \alpha_2\xi^2 + \alpha_3\xi^3$$

where

$$\xi = \frac{\phi}{(1-\phi)}$$
$$\alpha_1 = R_c^3 + 3R_c^2 + 3R_c + 1.5L_c(R_c^2 + 2R_c + 1)$$
$$\alpha_2 = 1.5(2R_c^3 + 3R_c^2) + 4.5L_c(R_c^2 + R_c)$$
$$\alpha_3 = 3R_c^3 + 4.5L_cR_c^2$$

When $L_c = 0$ we will have the result for the free energy change in transferring a spherical particle into the fluid. After calculating for dimer and monomer states, the change in free energy is calculated as,

$$\Delta\Delta F^{\phi} = \Delta F^{\phi}_{dimer} - 2 \times \Delta F^{\phi}_{monomer}$$

Instead of hard particle radius of crowder, the soft effective radius is used for SPT calculations. From the virial coefficient we have,

$$B = -2\pi \int e^{-\frac{\Phi(r)}{k_B T}-1} d^3r$$

Comparing with result for hard spheres,

$$B = \frac{2\pi\sigma^3}{3}$$

The effective crowder radius, $R_{effective} = \sigma$ - $R_p$.

Comparison of simulation with SPT calculation is shown in Fig. S5. The cylindrical length of dimer is chosen in order to conserve the volume of monomers. This is the reason for the observed higher stabilization from SPT prediction than simulation (Fig. S5 *orange circles*).

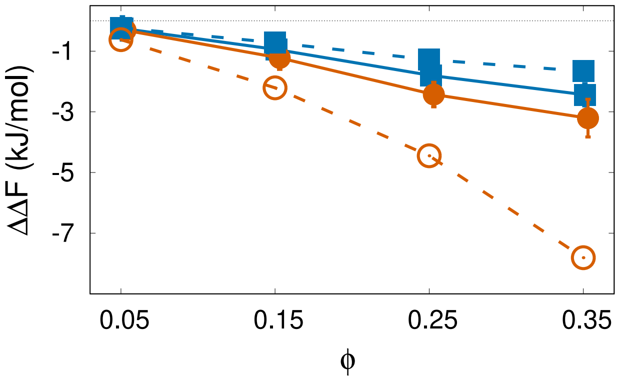


Figure S5: Free energy change in single crowder environments of hard sphere/repulsive 20 Å crowders (*blue squares*) and 12 Å crowders (*orange circles*). Solid, dashed lines indicate simulation and SPT prediction respectively. results 20 Scaled particle theory results (dashed lines) for corresponding simulations of single 20 Å (blue squares) and 12 Å (orange circles) crowders having harmonic repulsive interaction with protein.

### 4 Excluded volume calculation

The small (12 Å) crowders can entropically stabilize any of the pairs, i.e., 20 Å-20 Å crowder, protein-20 Å crowder or protein-protein. To investigate the entropic effect on various pairs, change in the volume accessible to the 12 Å crowder is calculated as follows.

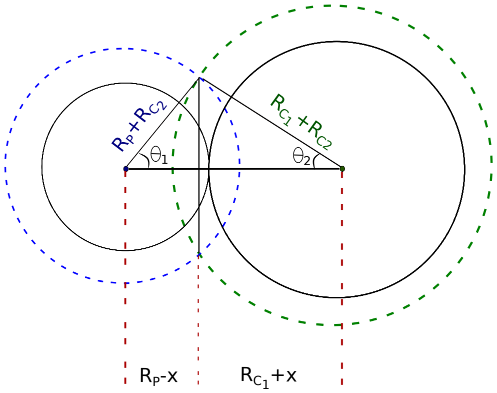


Figure S6: Overlap volume when protein and large crowder are in contact. The radius of solid circles is same as protein and large crowder size. The dashed circles represent the distance within which the center of the smaller crowder can not enter.

The volume of the lens (V) formed in the middle by the green and blue dashed spheres is equal to the sum of the volume of spherical caps. For two spheres of radii $R_p + R_{C_2}$ and $R_{C_1} + R_{C_2}$ with centers at a distance d = $(R_p + R_{C_1})$ apart, V can be calculated by geometric considerations and is given by,

$$V = \frac{\pi(R_p + R_{C_1} - d)^2(d^2 + 2dR_{C_1} - 3R_{C_1}^2 + 2dR_p + 6R_{C_1}R_p - 3R_p^2)}{12d}$$

As shown in Table. S2, the overlapping volume increases with increase in size of the species. It is entropically more favorable for small crowders to bring two species which has more overlap volume. Bringing two 20 Å crowders is most favorable, but it does not affect the stabilization of protein dimer directly. Bringing protein and 20 Å crowder is more favorable than bringing two proteins together, which directly affects the stability and reduces the overall stabilization of protein dimer. Fig. S7 shows different pairs which are stabilized in presence of small repulsive crowders.

Table S2: Volume excluded for 12 Å ($C_2$) crowder for different configurations of protein monomers and 20 Å ($C_1$) crowder. All volume units in $nm^3$ or $\times 1000 Å^3$

| Configuration | Overlap volume |
|---|---|
| Protein (13.5 Å)-protein (13.5 Å) | 19.45 |
| Protein (13.5 Å)-$C_1$ (20 Å) | 21.82 |
| $C_1$ (20 Å)-$C_1$ (20 Å) | 25.33 |

In the mixed crowder environment of only repulsive crowders at mixing ratio 1:3, presence of large number of smaller crowders increases the population of protein-large (20 Å) crowder configuration (Fig. S8 blue bars) which

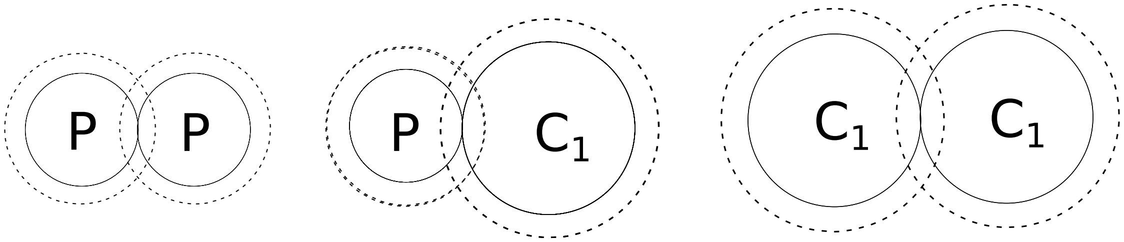


Figure S7: Overlap volume for protein-protein (P-P), protein-large crowder (P-$C_1$) and large crowder -large crowder ($C_1$-$C_1$) configurations.

decreases the stability of protein dimerization (Fig. S8 red bars) compared to only single crowder environment of smaller crowders (Fig. S8 gray bars). In mixed crowder environment, the entropic stabilization of protein-20 Å crowder pairs reduces stabilization of dimer (Fig. S8 *red bars*) w.r.t single crowder environment of 12 Å crowders (Fig. S8 *gray bars*).

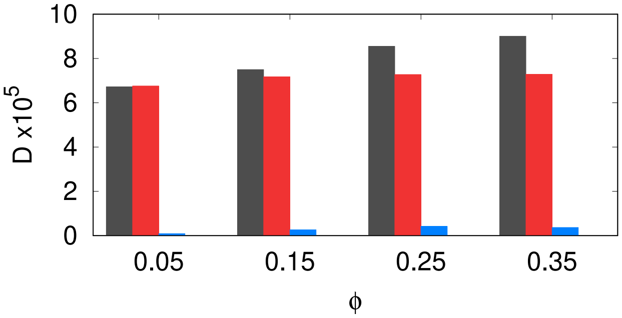


Figure S8: Dimer frames in single 12 Å crowder environment (*gray bars*), dimer frames for mixed repulsive crowders in 1:3 ratio (*red bars*), and protein-20 Å crowder population in the same mixed environment (*blue bars*).

#### 5 $\Delta\Delta H$ Calculation

Let the change in enthalpy change during association in crowded medium w.r.t no-crowder medium be given by,

$$\Delta\Delta \mathrm{H} = \Delta \mathrm{H}^{\phi} - \Delta \mathrm{H}^{0}$$

As there are no crowders at $\phi = 0$, we get $\Delta \mathrm{H}^0 = 0$. At crowder volume fraction $\phi$, we get the change in enthalpy of association as,

$$\Delta \mathrm{H}^{\phi} = \mathrm{H}^{\phi}_{dimer} - \mathrm{H}^{\phi}_{monomer}$$

.

From the simulation we obtain the average enthalpy in any state (dimer or monomer) as the sum of potential energy over all crowders ($C_1$ and $C_2$) over two GB1 proteins over all the frames averaged over all the frames. Mathematically it can be written as,

$$\mathrm{H}^{\phi}_{\mathrm{dimer}} = \frac{1}{\text{dimer frames}} \sum_{\text{frames}} \sum_{\text{GB1}} \sum_{C_1} \sum_{C_2} \mathrm{V}_{\mathrm{LJ}}|_{\mathrm{dimer}}$$

and

$$\mathrm{H}^{\phi}_{\mathrm{monomer}} = \frac{1}{\text{monomer frames}} \sum_{\text{frames}} \sum_{\text{GB1}} \sum_{C_1} \sum_{C_2} \mathrm{V}_{\mathrm{LJ}}|_{\mathrm{monomer}}$$

Enthalpy change from the average of histogram of enthalpy is given by,

$$\Delta\Delta \mathrm{H} = \Delta \mathrm{H}^{\phi}$$

$$= \mathrm{H}^{\phi}_{\mathrm{dimer}} - \mathrm{H}^{\phi}_{\mathrm{monomer}}$$

$$= \left[\frac{1}{\text{dimer frames}} \sum_{\text{frames}} \sum_{\text{GB1}} \sum_{C_1} \sum_{C_2} [\mathrm{V}_{LJ}|_{dimer}]\right] - \left[\frac{1}{\text{monomer frames}} \sum_{\text{frames}} \sum_{\text{GB1}} \sum_{C_1} \sum_{C_2} [\mathrm{V}_{LJ}|_{monomer}]\right]$$

$\Delta\Delta$H is then averaged over the eight parallel simulations for a particular $\phi$.

$-\mathrm{T}\Delta\Delta\mathrm{S}$ is calculated from the difference of change in free energy and change in enthalpy by the following relation,

$$-\mathrm{T}\Delta\Delta \mathrm{S} = \Delta\Delta \mathrm{F} - \Delta\Delta \mathrm{H}$$

#### 6 Single crowder environment with strong protein-12 Å LJ interaction ($\epsilon = 5$ kJ/mol)

In presence of strong protein-small crowder attractive strength ($\epsilon = 5$ kJ/mol), $D_{ex}^{in}$ increases with $\phi$ (Fig. S9b *blue bars*). The outside monomers within cutoff of different 12 Å crowders increases ($D_{ex}^{out}$ -ve, Fig. S9b *red bars*). Here, the protein-12 Å crowder transient units are treated as repulsive partners by other 12 Å crowders and thus it is entropically favorable to bring these units together. But the enthalpic cost hinders the contact formation between protein monomers and thus inhibits dimerization. The overall effect is destabilization (Fig. S9b *gray bars*).

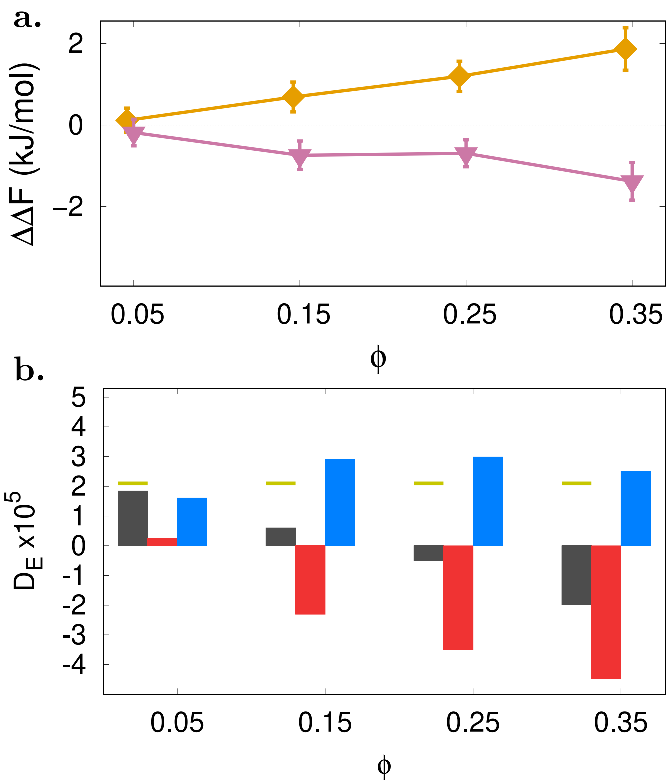


Figure S9: (a) Change in free energy in presence of single crowder environment with protein-12 Å crowder attraction strength 1 kJ/mol (*pink down-triangles*) and 5 kJ/mol (*yellow diamonds*). (b) For 5 kJ/mol LJ attraction strength, at each volume fraction, the bars are shown in the order of $D_{ex}$ (*gray bars*), $D_{ex}^{out}$ (*red bars*), and $D_{ex}^{in}$ (*blue bars*). The *yellow* line marking shows $D_{ex}$ for $\phi = 0$ case.

#### 7 Mixed crowder simulation with strongly attractive small (12 Å) crowders ($\epsilon = 5$ kJ/mol)

**Case V: HR-LJ ($\epsilon$ - $\epsilon$) interaction between protein and crowders**

In an environment in which the protein-smaller (12 Å) crowders have strong attractive strength and protein-larger crowder (20 Å) have repulsive interaction, Fig. S10a shows the overall change in free energy which also includes $\Delta\Delta F$ for single crowder species for reference. Fig. S10b and c shows the enthalpic and entropic contributions to $\Delta\Delta F$, respectively. Fig. S10d depicts possible monomer (F) and dimer (G) configurations. Fig. 10e and f plots the number of excess dimer configurations inside and outside of the cutoff distance of the smaller crowder at 3:1 and 1:3 crowder composition, respectively, providing an outlook on the enthalpic and entropic stabilization.

For 3:1 mixing ratio, dimer formation is stabilized for all volume fractions (Fig. S10a *olive up-triangles*). At this composition, it is entropically favored to bring protein monomers together by 20 Å repulsive crowders. Also, two or three small attractive crowders can transiently interact with the protein which acts as an unit (Fig. S10d). With increase in $\phi$, it is entropically favored to bring the protein monomers or transient protein-12 Å crowder units together (Fig. S10c *olive bars*) which can lead to formation of dimer. The corresponding monomer and dimer configurations are shown in Fig. S10d. This can also be seen with the increase in $D_{ex}^{in}$ with $\phi$ (Fig. S10e *blue bars*) showing increase in dimer configurations within the cutoff of 12 Å crowders and reduction in the dimer configuration forming outside the cutoff (Fig. S10e *red bars*). Hence, the overall stabilization is entropic at 3:1 mixing ratio. The reduced stabilization compared to single 20 Å crowder environment (Fig. 10a *blue squares*) can arise due to entropic stabilization in bringing protein-12Å transient unit and 20 Å crowder together.

At 1:3 composition, the 12 Å crowders are strongly interacting and occupies more volume than the larger crowder. With more 12 Å crowders, it is entropically more favored to bring the protein-12 Å transient units together. This leads to an increase in $D_{ex}^{in}$ inside the cutoff of 12 Å crowders and decrease in $D_{ex}^{out}$ (*blue* and *red* bars in Fig. S10f). At lower volume fractions, the entropic effect can stabilize both protein-12 Å transient unit and 20 Å crowder together or two protein-12 Å transient units together. At the highest $\phi$, entropic stabilization for the protein-12 Å crowder units, is highest.

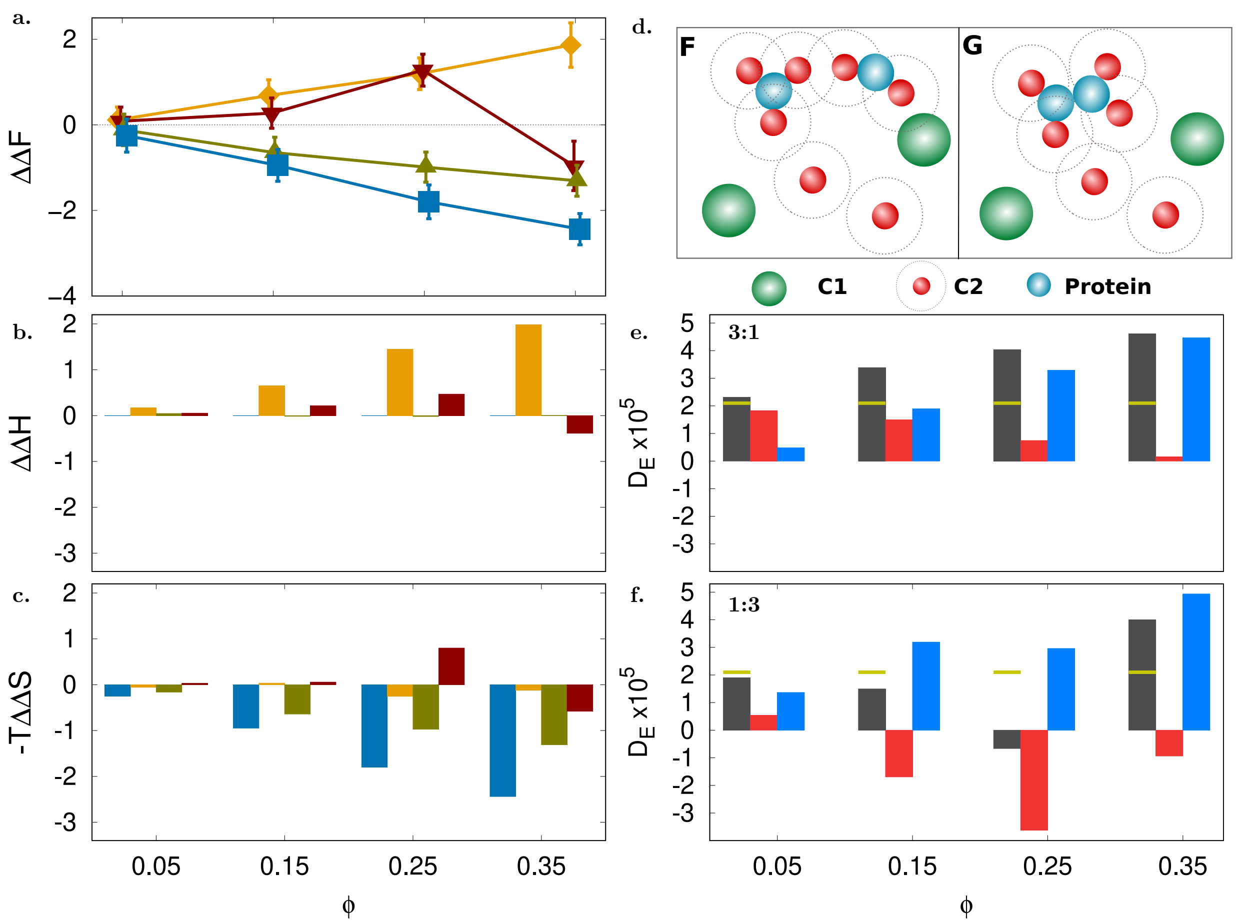


Figure S10: Dimerization in presence of mixed HR-LJ interactions. (a) Change in free energy $\Delta\Delta F$ in presence of a mixed environment of 20 and 12 Å crowders at mixing ratio 3:1 (*olive up-triangles*) and 1:3 (*red down-triangles*) interacting via HR and LJ potentials, respectively. $\Delta\Delta F$ for single crowder environments with protein-20 Å HR potential (*blue squares*), protein-12 Å LJ ($\epsilon = 5$ kJ/mol) potential (yellow diamonds) are shown for reference. (b) Bars showing change in enthalpy $\Delta\Delta H$ at each volume fraction $\phi$ in the order of single 20 Å, 12 Å crowders and mixed crowders in 3:1 and 1:3 ratios (bars follow the line colors defined in (a)). (c) Bars showing change in entropy $-T\Delta\Delta S$ at each volume fraction (as in (b)). (d) Major configurations contributing to the monomer and dimer frames. (e) Bars indicating excess dimers for 3:1 mixing ratio shown in the order of $D_{ex}$ (*gray*), $D_{ex}^{out}$ (*red*), and $D_{ex}^{in}$ (*blue*). The *yellow* line marks $D_{ex}$ for $\phi = 0$ case. (f) Same quantities as (e) for 1:3 mixing ratio. All energy units are in kJ/mol.

**Case VI: LJ-LJ ($\epsilon$ - $\epsilon$) strong interaction between protein and crowders**

Finally, consider both crowders having strong attractive LJ interaction with the protein. The protein-20 Å crowder has attraction strength of 8 kJ/mol, while the 12 Å crowder has a strength of 5 kJ/mol, mimicking small cellular components with strong 'sticky' interactions. This crowding scenario explores dimerization in an environment of competitive non-specific binders. Again, Fig. S11a-f describes similar quantities as in the previous case. As the large crowder is attractive, $D_{ex}$ $D_{ex}^{in}$ , and $D_{ex}^{out}$ are calculated w.r.t cutoff of 20 Å crowder.

For 3:1 mixing ratio, dimer formation stability closely follows that of the single 20 Å crowder (Fig. S11a *olive up-triangles*). At this composition, with increase in $\phi$, it is enthalpically favored (Fig. S11b *olive bars*) for dimer to form inside the cutoff of 20 Å crowder. This can be seen with increase in $D_{ex}^{in}$ with $\phi$ (Fig. S11e *blue bars*). It is also entropically favored to bring the protein-12 Å crowder transient units and 20 Å crowder together, resulting in entropic destabilization (Fig. S11c *olive bars*) of protein dimer and reduction of excess dimer frames ($D_{ex}^{out}$ ) outside the cutoff of 20 Å crowders (Fig. S11e *red bars*).

At 1:3 composition, the strongly interacting 12 Å crowders occupy more volume than 20 Å attractive crowders resulting in less (more) enthalpic stabilization (destabilization) (Fig. S11b *red bars*) due to 20 Å crowder than in 3:1 composition. This can be seen by increase in $D_{ex}^{in}$ (Fig. S11f *blue bars*) which is significantly less in magnitude compared to 3:1 composition. At lower volume fractions, there is competition between bringing protein-12 Å transient unit and 20 Å crowder together or bringing two protein-12 Å transient units together. This decreases the $D_{ex}^{out}$ upto $\phi$ = 0.25. At the highest volume fraction, the entropic stabilization effect is highest due to large number of small crowders which stabilizes the biggest units, i.e protein-12 Å transient units together and forming protein dimer resulting in reduced destabilization.

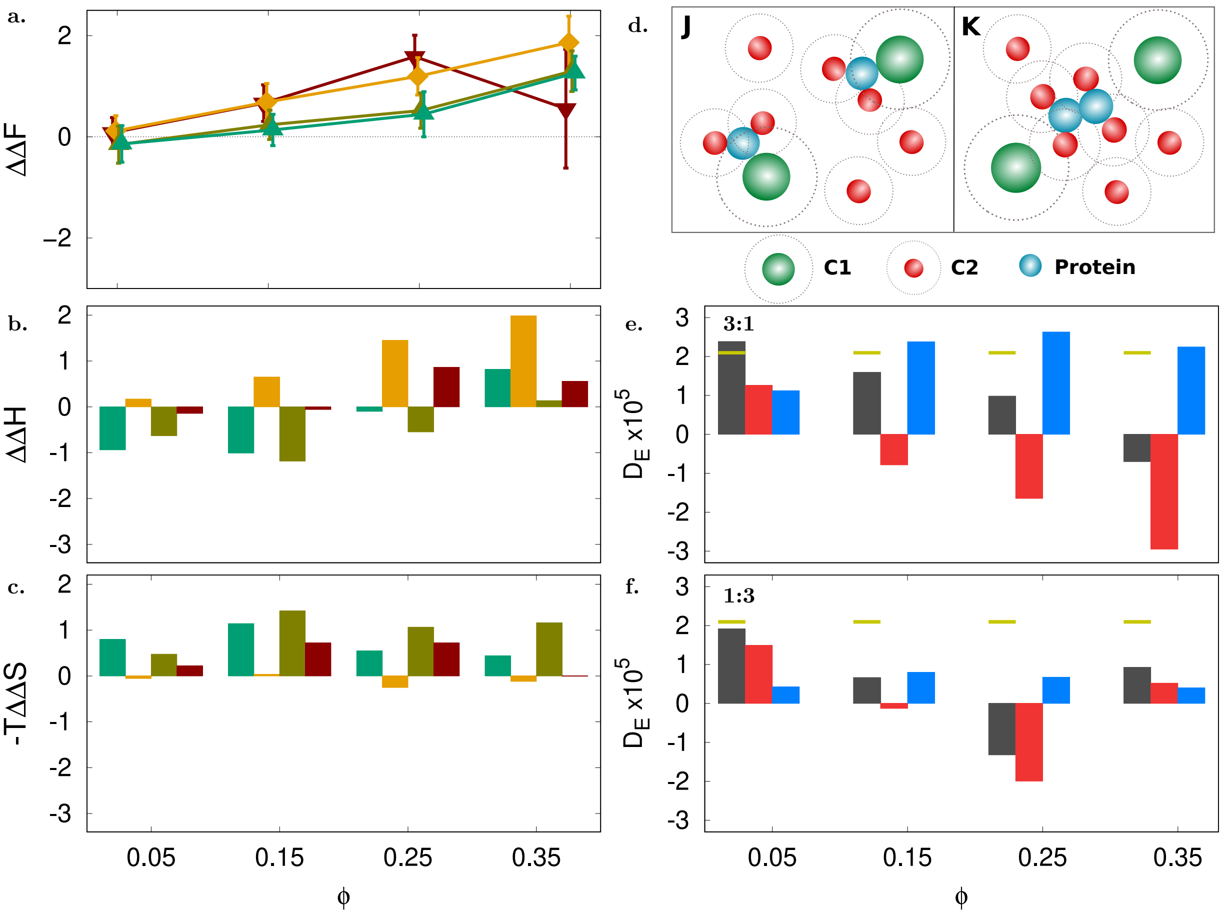


Figure S11: Dimerization in presence of mixed LJ-LJ interactions. (a) Change in free energy $\Delta\Delta F$ in presence of a mixed environment of 20 and 12 Å crowders at mixing ratio 3:1 (*olive up-triangles*) and 1:3 (*red down-triangles*) interacting via LJ and LJ potentials, respectively. $\Delta\Delta F$ for single crowder environments with protein-20 Å LJ ($\epsilon =$ 8 kJ/mol) potential (*green up-triangles*), protein-12 Å LJ ($\epsilon = 5$ kJ/mol) potential (*yellow diamonds*) are shown for reference. (b) Bars showing change in enthalpy $\Delta\Delta H$ at each volume fraction $\phi$ in the order of single 20 Å, 12 Å crowders and mixed crowders in 3:1 and 1:3 ratios (bars follow the line colors defined in (a)). (c) Bars showing change in entropy $-T\Delta\Delta S$ at each volume fraction (as in (b)). (d) Major configurations contributing to the monomer and dimer frames. (e) Bars indicating excess dimers for 3:1 mixing ratio shown in the order of $D_{ex}$ (*gray*), $D_{ex}^{out}$ (*red*), and $D_{ex}^{in}$ (*blue*). The *yellow* line marks $D_{ex}$ for $\phi = 0$ case. (f) Same quantities as (e) for 1:3 mixing ratio. All energy units are in kJ/mol.
